# A General Method to Accurately Count Molecular Complexes and Determine the Degree of Labelling in Cells Using Protein Tags

**DOI:** 10.1101/2023.09.08.556847

**Authors:** Stanimir Asenov Tashev, Jonas Euchner, Klaus Yserentant, Siegfried Hänselmann, Felix Hild, Wioleta Chmielewicz, Johan Hummert, Florian Schwörer, Nikolaos Tsopoulidis, Stefan Germer, Zoe Saßmannshausen, Oliver T. Fackler, Ursula Klingmüller, Dirk-Peter Herten

## Abstract

Determining the label to target ratio, also known as degree of labelling (DOL), is crucial for quantitative fluorescence microscopy and a high DOL with minimal unspecific labelling is beneficial for fluorescence microscopy in general. Yet, robust, versatile, and easy-to-use tools for measuring cell-specific labelling efficiencies are not available. This study presents a novel DOL determination technique named Protein-tag DOL (ProDOL), which enables fast DOL measurements and optimisation of protein-tag labelling. With ProDOL various factors affecting labelling efficiency, including substrate type, incubation time, and concentration, as well as sample fixation and cell type can be easily assessed. We applied ProDOL to investigate how HIV-1 pathogenesis factor Nef modulates CD4 T cell activation measuring total and activated copy numbers of the adaptor protein SLP-76 in signalling microclusters. ProDOL proved to be a versatile and robust tool for labelling calibration, enabling determination of labelling efficiencies, optimisation of strategies, and quantification of protein stoichiometry.

## Introduction

Fluorescence microscopy has long been a vital tool in biological research, enabling the detection of proteins of interest (POIs) in a variety of contexts. However, measuring copy numbers of a POI reliably with fluorescence microscopy not only requires a methodology to count fluorophores but also the information about the fraction of POI labelled with fluorescent markers. Labelling techniques such as immunolabelling result in variable and difficult to characterise labelling efficiencies. On the other hand, while genetic fusion with fluorescent proteins can yield a one-to-one ratio of label to the POI, it is often not suited for quantitative measurements and can be challenging due to insufficient photostability and ill-defined brightness states^1^. Additionally, fluorescent proteins in fusion constructs can exhibit a lower apparent labelling efficiency due to variable maturation efficiencies, or inefficient chromophore formation in different subcellular environments^2-6^. Self-labelling protein tags, such as SNAP-tag^7,8^ and HaloTag^9^ that bind at most one label per tag are also genetically fused to the POI and can bind a variety of fluorescent substrates with potentially superior photophysical properties^1^. However, due to the additional labelling step that requires incubation with fluorescent substrates, labelling efficiencies can vary depending on the chosen labelling condition. Moreover, unspecific binding of these substrates in the sample needs to be accounted for.

To ensure optimal labelling conditions for any labelling technique, the degree of labelling (DOL), or the ratio of fluorescent markers to POI, needs to be determined. A precisely determined DOL can also serve as a correction factor for measured protein copy numbers in complexes, and protein concentrations obtained from fluorescence microscopy techniques. However, determining the DOL can be challenging^10-12^ and different methods to address this issue have previously been developed^13^. One common approach is based on molecular counting standards such as the nuclear pore complex (NPC) combined with fluorophore counting methods such as super-resolution-based effective labelling efficiency (ELE)^10^, quick photobleaching step analysis (quickPBSA)^1^, or counting by photon statistics (CoPS) ^14,15^. Unfortunately, using protein complexes with known stoichiometry often comes with significant limitations, as complete complex assembly must be ensured and use of a homozygous knock-in cell-line is required. Additionally, methods such as ELE and CoPS require the use of labels suitable for super-resolution microscopy or specialised instrumentation, further limiting general application.

Colocalisation analysis with an additional, spectrally different label in close spatial proximity has been proposed previously, using e.g., a fusion between SNAP-tag and HaloTag to estimate the DOL of both tags^16^. Other work has utilised an additional antibody labelling against the protein tags^17^. However, all proposed methods can suffer from unspecific labelling of the reference signal resulting in an underestimation of DOL.

To overcome these limitations, we propose a modular DOL calibration probe that employs a fluorescent protein as a nearly background-free reference signal combined with protein tags. This construct can be transiently or stably expressed in various cell lines and provides a way to measure labelling efficiency through colocalisation at the single-molecule level, thus enhancing the reliability and versatility of the measurements. In the current implementation the DOL calibration probe is composed of a membrane anchored eGFP fused to SNAP-tag and HaloTag, and is named “protein-tag degree of labelling (ProDOL) probe”. Additionally, we developed a ProDOL analysis pipeline for labelling efficiency measurements by single-molecule colocalisation analysis. The integration of these techniques greatly enhances our ability to characterise fluorescent labelling approaches, measure labelling efficiencies, and thereby facilitate accurate counting of POIs. By identifying optimal labelling conditions for robust protein counting, we can better harness the potential of fluorescence microscopy in biological research, leading to a more accurate and reliable understanding of cellular processes. We demonstrate this potential by determining the time-resolved protein copy number of SLP-76 and phospho-SLP-76 in microclusters (MC) upon T cell activation, as well as how the cluster stoichiometry and T cell activation are affected by the HIV-1 protein Nef.

## Results

### Protein-tag degree of labelling – ProDOL

ProDOL is based on the colocalisation of the single-molecule signals emitted by labelled protein-tags and reference labels in spectrally separated images. To compute the DOL, the fraction of reference label colocalised with a protein-tag signal is assessed (Fig. 1b). We implemented this strategy by creating a fusion construct to serve as ProDOL probe with the ability to assess the labelling efficiency of different protein-tags in mammalian cell lines (see Methods for Addgene ID). The ProDOL probe was engineered with enhanced green fluorescent protein (eGFP) serving as a reference marker, and two self-labelling protein tags (HaloTag, and SNAP-tag). An N-terminal Lyn-kinase-derived membrane anchor targets the ProDOL probe to the plasma membrane via post-translational modification to enable single-molecule imaging by TIRF microscopy (Fig. 1a). An α-helical linker was added between SNAP-tag and HaloTag to facilitate maturation and avoid misfolding of the fusion protein, and a C-terminal His-Tag allows affinity purification or immunolabelling (Fig. 1a).

**Fig. 1:**
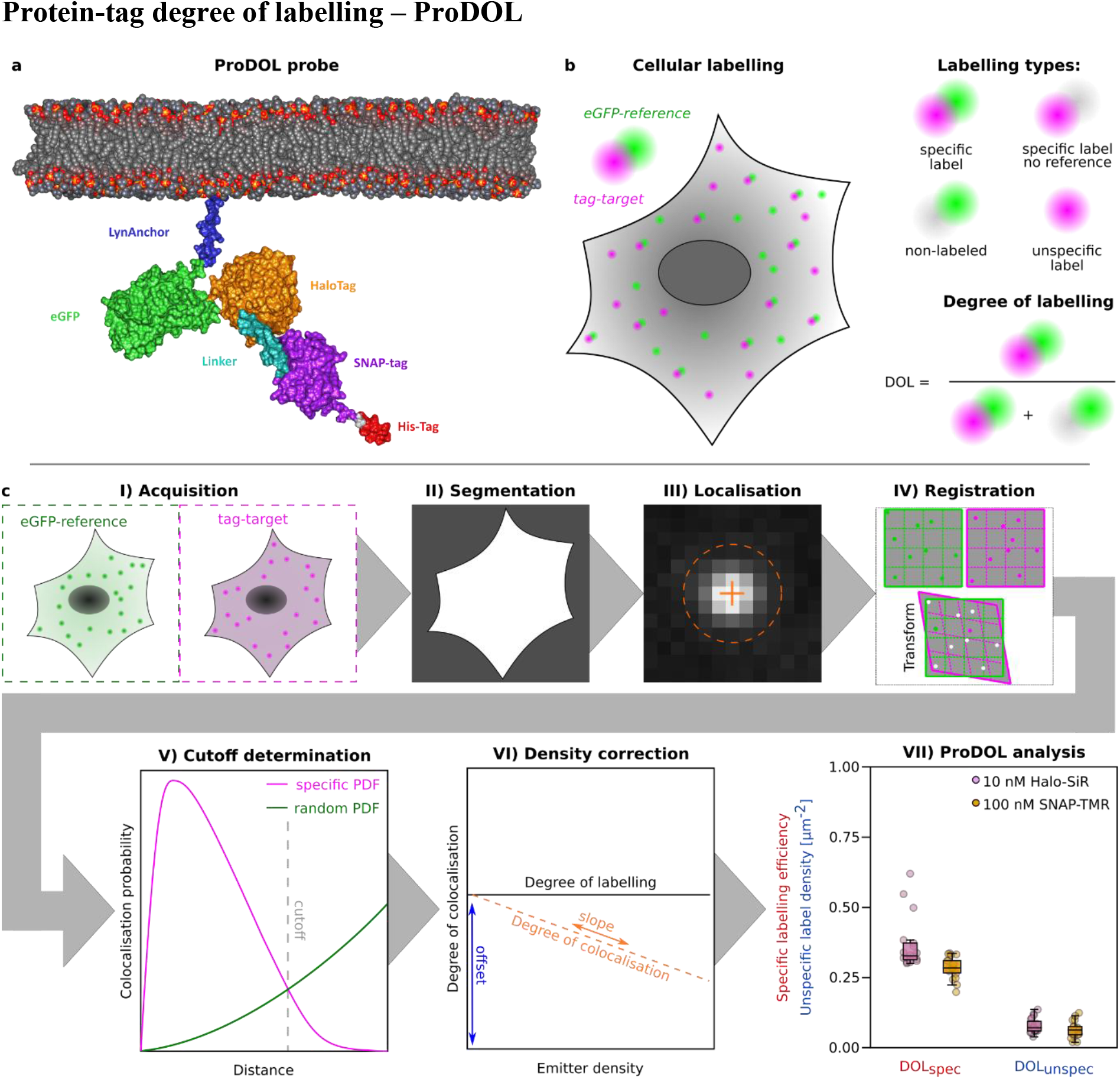
Principle and workflow of ProDOL. **a**, Molecular model of ProDOL probe generated using AlphaFold with post-prediction modification using published structures of individual components (PDB 2B3Q, 6Y8P, 6U32) and modelled lipid bilayer^35^. **b**, Schematic model to determine the degree of labelling based on single-molecule colocalisation analysis between an eGFP-reference and tag-target label. Within the ProDOL analysis four labelling types can be discerned (specific label, specific label with no reference, non-labelled and unspecific label) and used to calculate the degree of labelling and unspecific labelling density. **c**, ProDOL analysis workflow. **I**) Acquired images having both reference and target channels are used as input. **II**) Reference channel is used for generating a cell mask. **III**) Using SMLM fitting, sub-pixel location of reference and target are determined. **IV**) The channels are aligned using affine registration of the localisation data. **V**) Calculation of colocalisation cut-off T. **VI**) Factors are determined to correct the DOL for emitter density. **VII**) The DOL is determined for all acquired tag-target channels.

With eGFP as reference label the ProDOL probe enables spectral discrimination of frequently used red and far-red fluorescent SNAP-tag and HaloTag substrates. If required, eGFP can be replaced by alternative fluorescent proteins to facilitate DOL measurements for protein-tag substrates with excitation or emission spectra overlapping with eGFP. In a similar fashion, alternative protein tags can be introduced into ProDOL probe.

The expression level of the ProDOL probe is a critical parameter for robust DOL measurements. While high probe density provides better statistics, the density must be sufficiently low to reliably detect individual signals in diffraction-limited images. Therefore, the ProDOL probe was inserted into a retroviral pBABE plasmid to facilitate stable genomic integration and to achieve expression levels suitable for localisation of diffraction-limited single-molecule signals^18^.

Unspecific background signal is another important factor interfering with the determination of the DOL in living cells. While eGFP provides a background-free label, unspecific background has been frequently observed when labelling SNAP-tag or HaloTag in cells^19^. Therefore, we also created a truncated version of the ProDOL probe containing only the Lyn kinase membrane anchor and the eGFP reference label, but no additional protein tags (Lyn-eGFP – LynG). The LynG probe serves as a control to enable monitoring the density of unspecific labelling (Supplementary Fig. 1).

To complement the probe, we created a data-processing workflow consisting of seven modules accounting for the signal localisation in cells, chromatic aberrations, unspecific signals and variations of the label densities (Fig. 1c). First, colour multiplexed single-molecule images are acquired by TIRF microscopy (I), and a segmentation mask is generated from the reference channel image to exclude background signals outside of cells from subsequent analysis (II). The reference and label signals are localised with sub-pixel accuracy using ThunderSTORM^20^ (III) and subsequently corrected for chromatic aberrations by applying an affine transformation matrix (IV). Next, the colocalisation of the labelled tags with the reference is determined (V) using a distance cut-off *T* at which the fraction of specific colocalisation is maximised while the contribution of random colocalisation is kept at a minimum. The colocalised signals are then adjusted to account for effects of the probe density on signal detection (VI, see Supplementary Fig. 2). Multiple factors such as overlapping localisations, missed localisations, multiple assignment of signals, or cut-off *T* depend on the emitter density and can lead to an underestimation of the DOL. To obtain a robust result, DOL values determined in individual cells are averaged (VII).

**Fig. 2.**
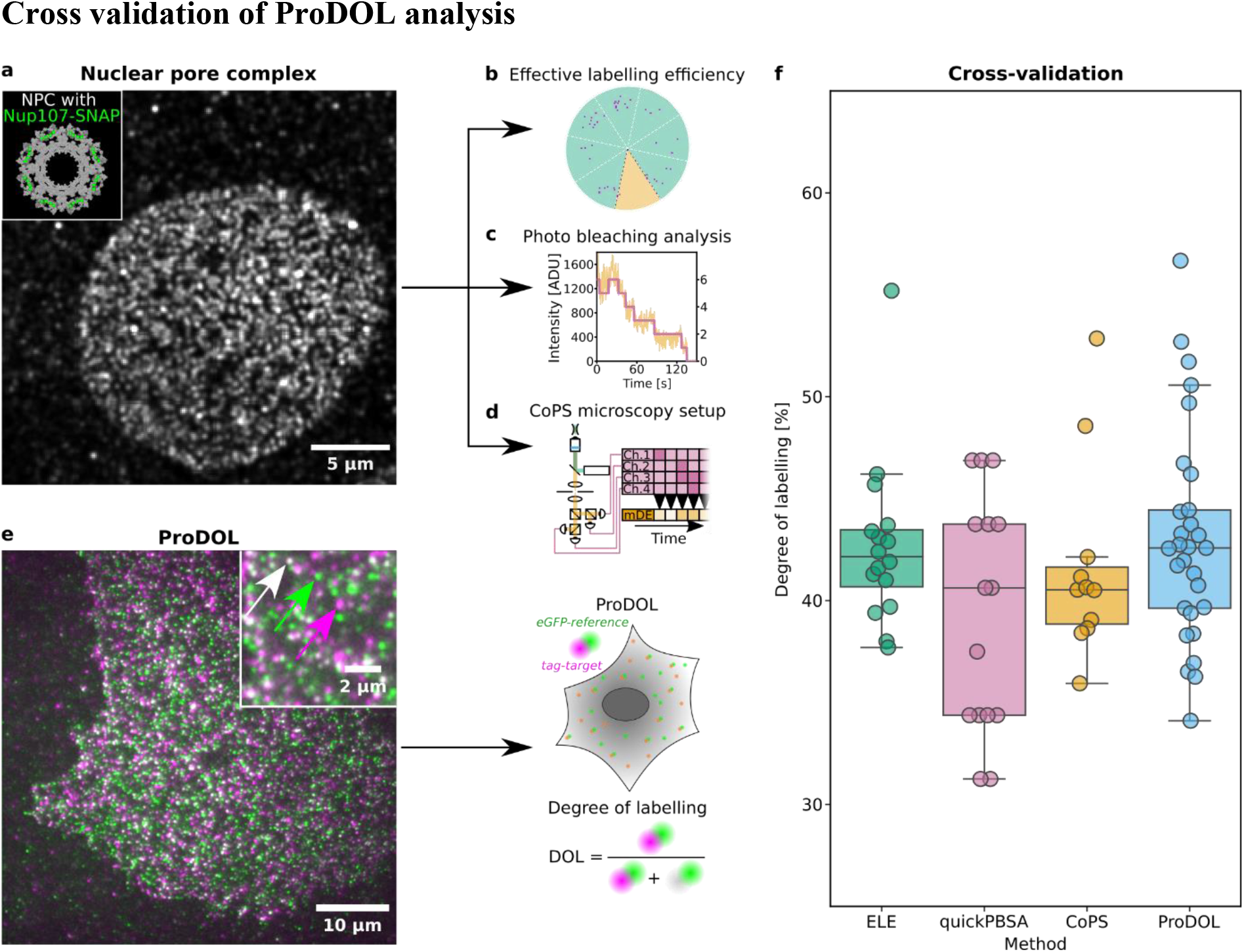
Comparison of molecular quantification techniques and ProDOL. **a**, Representative TIRF image of SNAP-AF647 stained U2OS cell with Nup107-SNAP knock-in. Inset: Structural model of NPC highlighting the expected distribution of the 32 Nup107 (green) copies across the 8 corners of NPCs. **b**, ELE analyses the distribution of single-molecule localisations across individual NPCs and fits a binomial function to the number of occupied corners to estimate the degree of labelling. **c**, PBSA relies on the stepwise reduction in fluorescence intensity (yellow) caused by photo bleaching. Here, Bayesian statistics are used to fit a change of states which correspond to the remaining emitting fluorophores (purple). **d**, CoPS microscopy setup with four independent Single-Photon Avalanche Detectors (SPADs) and a pulsed laser. A Time-Correlated Single Photon Counting System (TCSPCS) is used to generate a time trace of multiple detection events (mDEs). The histogram of which is matched to a probability model to determine the number of emitters. **e**, TIRF image of a U2-OS cell transiently transfected with the ProDOL probe. Arrows depict single-molecule localisations for eGFP-reference signal (green), SNAP-AF647 (magenta), and colocalisation of both (white). **f**, Box and whisker plot of cell-wise DOLs determined by the indicated methods. The data is grouped in a cell-to-cell basis. 73 cells were analysed (ELE: 16, quickPBSA: 17, CoPS: 11, ProDOL: 29). Label distribution per NPC can be found in Supplementary Fig. 6.

### Cross validation of ProDOL analysis

To experimentally validate the ProDOL method, we compared it to three approaches relying on defined protein complexes and different protein counting approaches. As calibration target for these experiments, we used nucleoporin 107 (Nup107), a key component of the nuclear pore complex (NPC), present at 32 copies per complex^21^. Nup107-SNAP was expressed in a genome-edited U2OS cell line from its native locus and was labelled with SNAP-Alexa Fluor 647 (SNAP-AF647)^22^ following a shared staining protocol for all counting techniques (Fig. 2a). For protein counting, we used three orthogonal emitter counting methods: effective labelling efficiency (ELE) ^10^, quickPBSA^1^, and CoPS^14,15^. ELE takes advantage of the known 8-fold symmetry of the NPC to infer the DOL from analysing single-molecule localisations within individual NPCs^9^ (Fig. 2b). quickPBSA detects stepwise intensity changes due to photobleaching during prolonged imaging to estimate the number of fluorophores^1^ (Fig. 2c). CoPS is a quantum imaging technique estimating the number of emitters in diffraction limited structures based on the detection of coincident photons^14^ (Fig. 2d). In contrast to ELE, CoPS and quickPBSA do not require a priori knowledge of the underlying geometry or a specific fluorescent label.

Finally, ProDOL measurements were conducted in transiently transfected U2OS cells expressing the ProDOL probe (Fig. 2e) using the same labelling conditions as for Nup107-SNAP described above. ProDOL analysis of the achieved labelling efficiency under these conditions revealed a DOL of 42.5±5.3%. This is in excellent agreement with the alternative quantification methods relying on the conserved copy number of Nup107/NPC where we determined labelling efficiencies of 42.2±4.1%, 37.5±8.0% and 40.5±4.9% for ELE, quickPBSA, and CoPS, respectively (Fig. 2f). Overall, a high degree of agreement across all four tested methods without significant differences (p=0.135, Kruskal-Wallis) between approaches demonstrates that ProDOL provides reliable DOL estimates without any specific requirements concerning the cell line and the fluorescent label and without requiring sophisticated emitter counting methods.

### Measuring the degree of specific and unspecific labelling

After successful validation, we utilised the ProDOL approach as a rapid screening tool to optimise labelling conditions by systematic variation of experimental parameters such as substrate concentration and incubation time. Optimal conditions for both pre- and post-fixation labelling with SNAP-SiR and Halo-TMR in HeLa cells were determined by inspection of DOL and unspecific labelling, with expression of ProDOL probe and LynG respectively (Fig. 3a). As expected, a steady increase of the DOL for both protein tags with increasing substrate concentrations (Fig. 3a I and III) both pre-(green) and post-fixation (yellow) was observed showing saturation at substrate concentrations above 100 nM. When labelling pre-fixation i.e., in living cells, both substrates plateau before 40%, with the Halo-TMR reaching 35±8% and the SNAP-SiR reaching 27±4%. In comparison, post-fixation labelling with SNAP-SiR showed a considerable increase in DOL (58±7%), while the Halo-TMR was only slightly higher at 40±8%. In addition, the DOL achieved by the protein tags differ in their concentration dependence in agreement with previously reported ligand affinities^23^. While labelling of HaloTag can be achieved at sub-nanomolar substrate concentrations, we could only detect specific binding for SNAP-tag at concentrations above 1 nM. As this is the case, under both pre- and post-fixation labelling conditions, differences in the cell permeability of the dye substrates can be ruled out as reason for this behaviour.

**Fig. 3:**
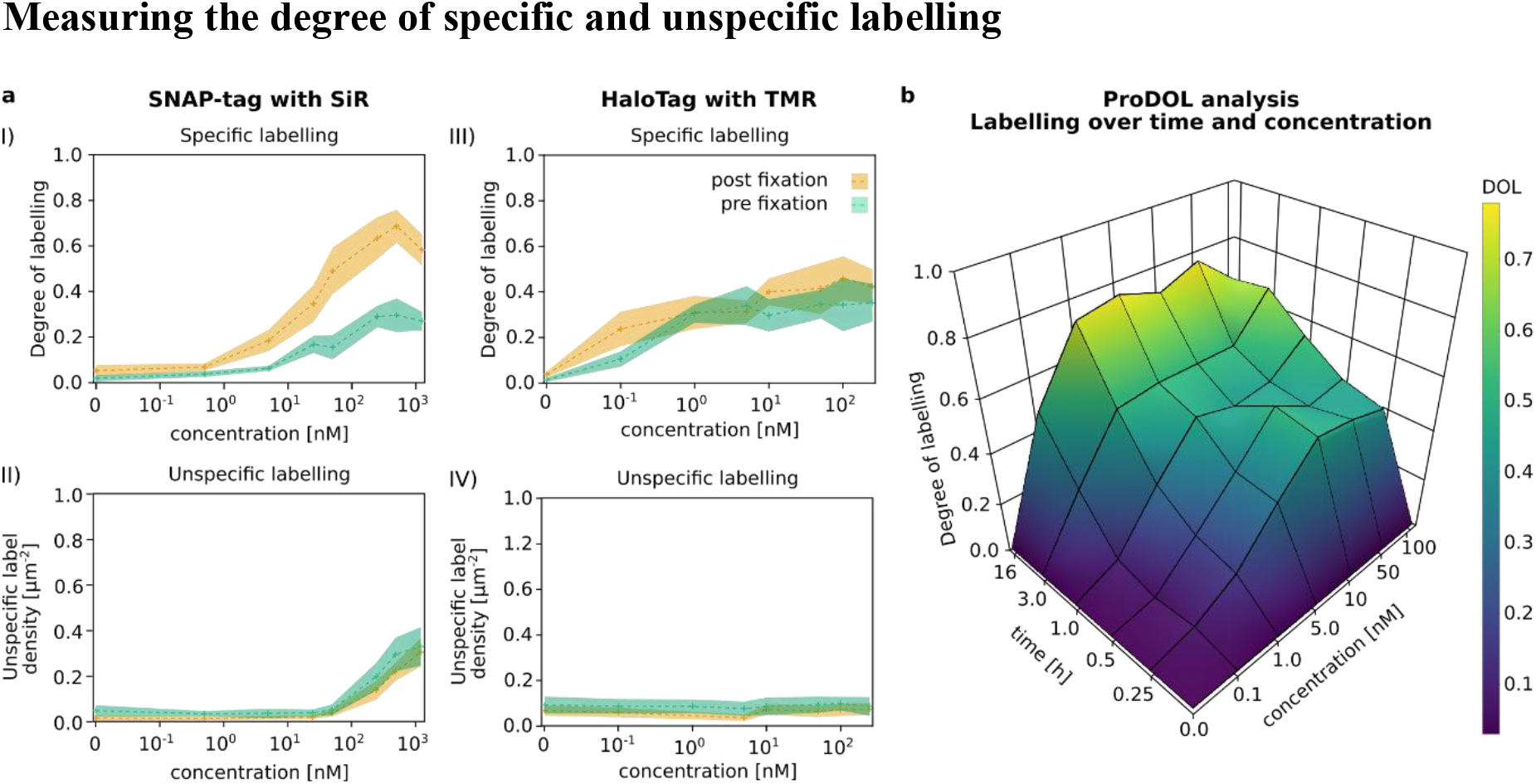
Determination of DOL as a parameter of incubation time, dye concentration and labelling condition. **a**, HeLa cells expressing ProDOL and LynG probes were stained with varying concentrations of SNAP-SiR (**I, II**) and Halo-TMR (**III, IV**) for 30 min pre-fixation (green) or post-fixation (orange). **I, III**) Determined DOL using ProDOL at varying substrate concentrations. **II, IV**) Unspecific labelling density measured in HeLa cells expressing LynG at varying substrate concentrations. **b**, DOL as function of incubation time and Halo-SiR concentration in Jurkat T cells. **a**, Median±SD (crosses, shaded bands), 6-20 cells per condition, **b**, Median, 20-38 cells per condition.

Knowledge of the DOL is essential for the quantification of protein copy numbers. However, it might not be sufficient if unspecific labelling reaches significant levels. To assess the density of unspecific labelling, the LynG probe was used in HeLa cells under identical labelling conditions as for the ProDOL probe and the unspecific labelling density was determined (Fig. 3a II and IV). While we observed little difference between pre- and post-fixation conditions, SNAP-SiR shows a significant increase in unspecific labelling beyond substrate concentrations of 100 nM, in contrast to Halo-TMR which shows less unspecific staining. Additional experiments with a range of different dye substrates in Huh-7.5 cells indicate that this is probably due to their molecular properties, such as polarity and charge (Supplementary Fig. 3). Moreover, we cannot exclude effects associated with the cell type as we observed significant variations in the achieved DOL across different cell lines labelled with the same dye substrate (Supplementary Fig. 4e).

**Fig. 4.**
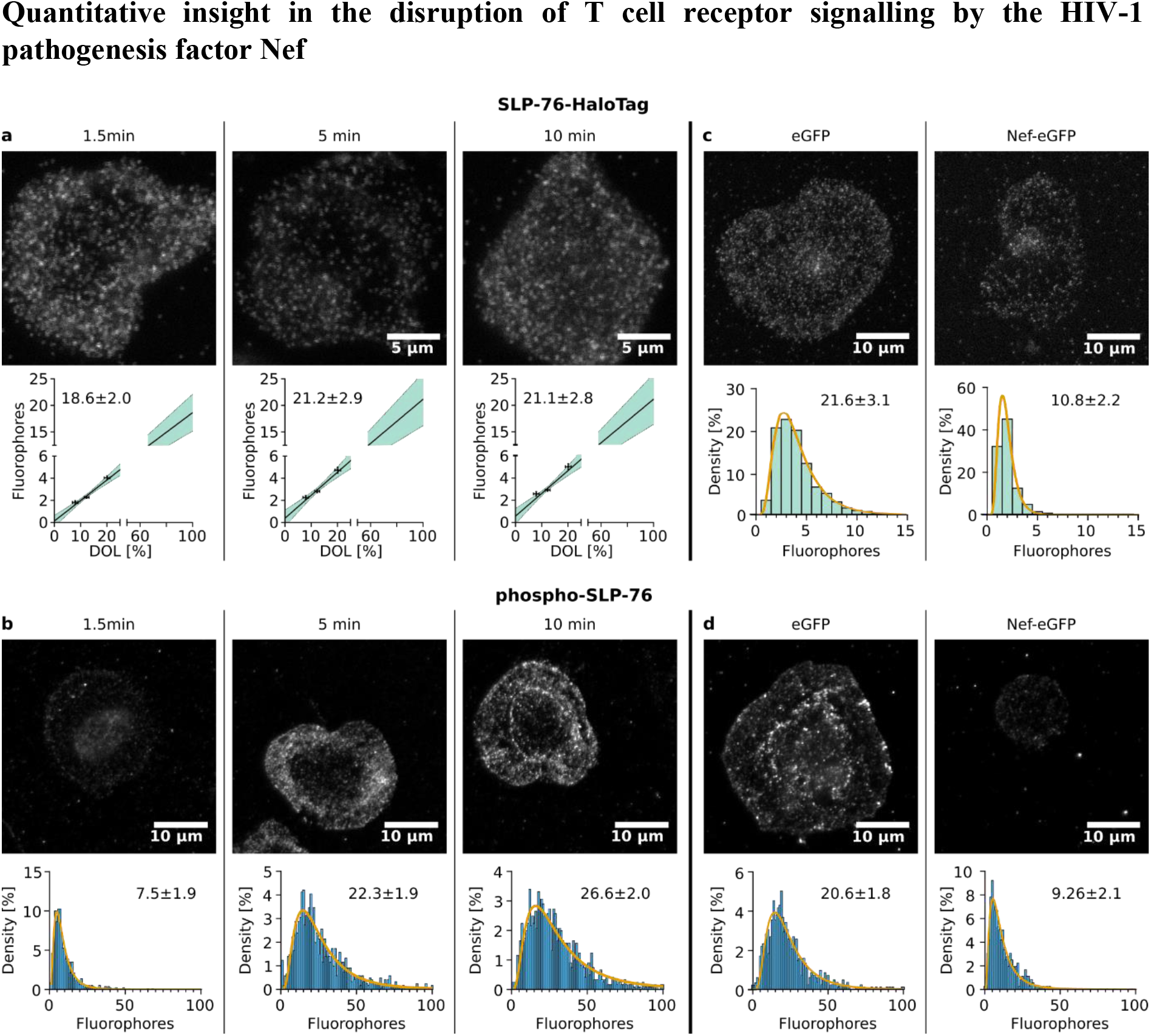
SLP-76 stoichiometry and phosphorylation inside activated Jurkat CD4 T cells. **a**,**c**, Jurkat CD4 T cells expressing SLP-76-HaloTag stained with Halo-SiR. **b**,Jurkat CD4 T cells labelled with anti-pY145-SLP-76-SiR. SLP-76 (**a**) and phospho-SLP-76 (**b**) clusters at different time points after activation (1.5 min, 5 min, 10 min). SLP-76 (**c**) and phospho-SLP-76 (**d**) clustering in Jurkat CD4 T cells transiently transfected with eGFP or Nef-eGFP. **a**, bottom panel: Extrapolation of the copy number of SLP-76-HaloTag per cluster using different degrees of labelling (11, 16, 27%). Shaded region: 95% CI. **b**, Absolute quantity of phospho-SLP-76 phosphorylation at different time points after activation (1.5 min, 5 min, 10 min). **c**, bottom panel: Histogram of labelled SLP-76-Halo per cluster (at DOL=16.7±1.7%) in the presence and absence of the viral protein Nef. **d**, Effects of Nef on SLP-76 phosphorylation. **a**-**d**, top panels show representative confocal images of phospho-SLP-76 stained with SiR. **b**-**d**, bottom panel: Histogram of SiR-labelled phospho-SLP-76 and SLP-76 copy numbers were fitted to a log-normal probability function (yellow). Log-normal fit: GeoMean ± GeoSD. Linear regression: Mean ± SE. **a**, DOL: 8-27 cells each, CoPS: 5-7 cells, 72≤N≤268 clusters, **b**, 20-27 cells 817≤N≤1268 clusters, **c**, 380≤N≤1348 clusters, **d**, 26-28 cells, 358≤N≤822 clusters.

To show the generalisability of the ProDOL approach for optimising labelling conditions, we performed further experiments with Jurkat T cells (Fig. 3b) and H383 cells expressing ProDOL labelled with Halo-TMR and SNAP-SiR (Supplementary Fig. 4a-d). As expected, the DOL increases with incubation time and substrate concentrations. In combination with the associated quantification of unspecific labelling (Supplementary Fig. 3c,d), this allows the selection of optimal labelling conditions for experiments quantifying protein copy numbers in cells by means of fluorescence microscopy.

### Quantitative insight in the disruption of T cell receptor signalling by the HIV-1 pathogenesis factor Nef

The HIV-1 pathogenesis factor Nef alters the activation of infected CD4 T cell in response to stimulation of the T cell receptor (TCR) to expand the life span of activated cells and thus optimise the production of viral progeny^24,25^. TCR engagement triggers the formation of signalling competent protein microclusters (MCs) whose composition and activity determines magnitude and breadth of signalling outputs. Our previous study suggested that Nef affects TCR signalling by reducing the formation of MCs that contain SLP-76, an adaptor protein and central coordinator of TCR signalling^26^, but we lack quantitative information on the protein composition and activity of individual MCs^27^. To address this, we first used ProDOL in combination with CoPS to determine the absolute SLP-76 copy number in response to TCR engagement in the absence of presence of Nef. To this end, Jurkat CD4 T cells stably expressing a SLP-76-HaloTag fusion protein were labelled in suspension with Halo-SiR and stimulated on glass cover slides functionalised with α-CD3 antibodies for activation^28-30^. To synchronise the activation reaction of individual cells, only cells that adhered to the cover slide within the first 3 seconds were observed and all other cells washed away. We selected single cells in which individual SLP-76 clusters labelled with Halo-SiR can be optically resolved as distinct spots in confocal fluorescence microscopy images upon settling on glass substrates for counting with CoPS (Fig. 4). SLP-76 MC formation during T cell activation was monitored by fixing the incubated samples at different times after activation (1.5, 5, and 10 min) for subsequent CoPS experiments. In parallel, the DOL was monitored in Jurkat T cells stably expressing the ProDOL probe to enable extrapolation of the labelled molecule numbers to absolute protein copy number of SLP-76-HaloTag in MCs (Fig. 4a). Surprisingly, our measurements indicate no significant difference (p=0.7495) between 1.5 and 10 min of activation, with the number of SLP-76-HaloTag remaining constant at around 20 protein copies per cluster. Interestingly, the associated label number distributions determined at these times show a log-normal distribution of increased variance (Fig. 4b-d) clearly indicating a lack of defined cluster organisation which can be explained by the multiplicative product of independent variables in accordance with the central limit theorem in log-space commonly found in intracellular distributions^31^.

Having established that the overall abundance of SLP-76 in MCs is relative constant following TCR engagement, we next determined the number of SLP-76 copies carrying an activating phosphorylation at tyrosine 145 with a phospho-specific anti-SLP-76 antibody (Fig. 4b). The antibody was labelled with SiR at a DOL of 1.27±0.05 as measured by absorption spectroscopy to approximate a 1:1 stoichiometry of label vs. phosphorylation site. Determining the label number distribution of the labelled antibodies using CoPS (Supplementary Fig. 5) enabled a more quantitative assessment of the phosphorylation during T cell activation. This revealed that SLP-76 phosphorylation significantly increased over time from 7.5±1.9 at 1.5 min to 22.3±1.9 at 5 min (p<0.0001), and to 26.6±2.0 at 10 min (p<0.001) (Fig. 4b). Using the dye to antibody ratio (DOL: 1.27±0.05) a corrected quantity of labelled phospho-SLP-76 per MC d, can be determined for 1.5 min: (5.9±1.5), 5 min: (17.6±1.6), and 10 minutes (20.9±1.8) accounting for the total population of both SLP-76 and SLP-76-HaloTag. Together, these results indicate that the function of SLP-76 in early TCR signalling is regulated by phosphorylation in individual MCs and not by recruitment of SLP-76 into MCs.

Finally, we assessed which of these parameters are affected by HIV-1 Nef and transiently expressed an HIV-1 Nef-eGFP fusion protein or an eGFP control in Jurkat CD4 T cells stably expressing SLP-76-HaloTag. In addition to the expected reduction in the overall number of SLP-76 MCs^27^, evaluating the stoichiometry of SLP-76 MCs revealed a significant reduction of SLP-76-HaloTag copies per MC 5 min post activation (eGFP: 21.6±3.1; Nef-eGFP 10.8±2.2, p<0.0001) (Fig. 4c). Consistently, the density of phospho-SLP-76 containing MCs (Supplementary Fig. 5c) and the amount of phospho-SLP-76 per MC was also significantly reduced in presence of Nef (eGFP: 9.26±2.1; Nef-eGFP: 20.6±1.8, p<0.0001; DOL corrected: eGFP: 7.3±1.7; Nef-eGFP: 16.2±1.6) (Fig. 4d). Notably, the presence of Nef did not affect the proportion of SLP-76 phosphorylation per MC (Nef-eGFP: 45±11% vs eGFP: 50±13% -358≤N≤1348) (p=0.0620). Together, the results define that Nef disrupt SLP-76 function at the level of MC recruitment but does not affect the activation of MC resident SLP-76. This accurate quantification of SLP-76 using ProDOL gives us an unprecedented look into the stoichiometry of protein complexes.

## Discussion

The ProDOL probe in combination with the established data analysis pipeline provides a robust and versatile labelling calibration workflow. Unlike other approaches ProDOL allows for determination of the degree of labelling across a variety of mammalian cell lines with potential use in other species where transient expression of a plasmid construct is possible. Additionally, ProDOL can be carried out on any single-molecule sensitive microscopy setup with diffraction-limited acquisition as no super-resolved acquisition is needed, expanding both on the feasibility, and dye compatibility. We have further validated the robustness of ProDOL in determining labelling efficiencies and its sensitivity to small methodological changes with extensive experiments and simulations (Supplementary Fig. 2). Thereby ProDOL can be used to rapidly and robustly determine optimal labelling conditions for protein-tag labelling and to measure absolute protein copy numbers in cells.

Determination of DOL assumes similar labelling efficiencies between the POI and the DOL probe. While this is not guaranteed, we provide evidence that similar DOLs are achieved for the plasma-membrane localised ProDOL probe and for NPCs. In addition, this also shows that comparable labelling efficiencies are achieved for protein tags sparsely distributed across the plasma membrane as well as for nucleoporins which are known to form high density structures in the nuclear membrane. However, additional subcellular compartments such as the lumen of cellular organelles or the extracellular leaflet of the plasma membrane warrant further studies to systematically dissect the influence of subcellular protein tag localisation on achieved DOL.

ProDOL highlights the need for robust and easy to implement labelling calibration, as most if not all labelling strategies results in sub-stoichiometric labelling^10^. Interestingly, PFA fixation increased the maximal labelling efficiency achieved for SNAP-tag labelling while HaloTag labelling was not significantly affected.

By applying the ProDOL concept to T cell activation, we showed how the composition of signalling MCs changes over time. We were able to demonstrate that the copy number of SLP-76 per MC stays constant between 1.5 and 10 min after activation, but phosphorylation increases over time. Thus, SLP-76 is either recruited in an early phase of T cell activation not covered by these experiments or already pre-assembled in MCs prior to activation. We also applied ProDOL to study how the HIV-1 pathogenesis factor Nef rewires TCR signalling to the benefit of the virus. Here, we established for SLP-76 that the viral protein primarily acts to prevent recruitment of host cell components into signalling hubs. However, successfully assembled signalling MCs are protected from manipulation by Nef and thus show similar phosphorylation ratios of SLP-76. ProDOL will now enable analogous analysis for more components of the TCR signalling machinery and profoundly impact the development of strategies to therapeutically restore physiological TCR signalling in HIV-1 infected CD4 T cells.

We expect that ProDOL will be implemented for a variety of uses, including but not limited to label optimisation, protein stoichiometry determination, and as quality control run along experiments for validating the labelling protocol. Optimising labelling strategies is especially useful for super-resolution imaging, as high labelling density with minimal background signal is crucial for appropriate sampling of structures at highest resolution. Having the ProDOL analysis available further enables the community to develop upon the concept of determining the DOL using colocalisation analysis. With the existing ProDOL probe, the labelling efficiency of direct immunolabelling can be determined when using anti-GFP or anti-His-Tag antibodies, commonly used due to the abundance of fusion constructs with these tags. Furthermore, ProDOL probe versions carrying any other peptide or protein tags such as CLIP, TMP-tag or HA-tag^32-34^ can easily be implemented by modifying the existing ProDOL probe. Moreover, the construct could be transferred into plasmids optimised for different origins opening up their use in non-mammalian cells lines such as plant cells or bacteria.

## Methods

### Cloning

A ProDOL precursor was synthesized as gBlock (IDT) consisting of Lyn kinase membrane anchor (N-terminal amino acid: 1-13), α-helical linker^36^, SNAP-tag (SNAP26m) and His-Tag. Between all segments unique restriction sites were included allowing for easy modification by restriction/ligation cloning (Supplementary Fig. 1). The precursor was inserted into pMOWS vector using EcoRI, PacI restriction sites ^37^.

eGFP and HaloTag were inserted between LynAnchor and linker using the unique restriction sites XhoI, BamHI and AgeI. Next, SNAP26m-tag was exchanged to SNAP_f_-tag using NdeI and MfeI restriction sites. Finally, the full ProDOL probe was cloned into the pBABE expression vector.

For generating the LynG construct, pBABE-ProDOL was digested using BamHI and MfeI, leaving only LynAnchor and eGFP. After blunting the ends, the plasmid was re-ligated generating pBABE-LynG. Successful integration was confirmed for each vector using restriction digestion.

For generating stable Jurkat CD4 T cell lines, ProDOL and LynG constructs were transferred into a pWPI vector. For this, both constructs were PCR amplified from the respective pBABE vector with addition of PmeI and SpeI restriction sites during PCR. PCR products were ligated into the pWPI-vector. Successful integration was confirmed for each vector using restriction digestion.

Plasmids available through Addgene (pBABE-ProDOL probe-ID: 206866; pBABE-LynG probe-ID: 206867).

### Cell Culture

Adherent mammalian cells were cultured in a complete growth medium consisting of phenol red-free Dulbecco’s Modified Eagle Medium (DMEM), supplemented with 10% Fetal Bovine Serum (FBS), 1x GlutaMAX, and 1 mM sodium pyruvate (all Gibco). Cells were incubated at 37 °C, 5% CO_2_, and 100% humidity. Adherent cells stably expressing ProDOL or LynG probes were additionally treated with 1.5 µg/mL puromycin to maintain transgene expression. Genetically modified cells were additionally treated with the appropriate selection antibiotics. Upon reaching approximately 80% confluency, cells were subcultured. The protocol involved aspiration of growth medium, a single rinse with PBS, and subsequent incubation with TrypLE Express until cells fully detached. Inactivation of TrypLE was achieved via the addition of twice the volume of complete growth medium, followed by centrifugation at 500× g for 5 min to pellet the cells. After removal of the supernatant, the cell pellet was resuspended in complete growth medium, and subsequently seeded at dilution ratios ranging from 1:6 to 1:10.

Suspension cells were cultured in complete growth medium consisting of phenol red-free RPMI-1640 medium, 10% FBS, 1x GlutaMAX, 1 mM sodium pyruvate, and 1x Penicillin-Streptomycin (Gibco). Cells were incubated at 37 °C, 5% CO2, and 100% humidity. Genetically modified cells were additionally selected by the appropriate selection antibiotics. All suspension cells were passaged into fresh medium to maintain a concentration of between 2× 10^5^ and 8× 10^5^ cells/ml.

### Stable cell line generation

Stable adherent transgenic cell lines, expressing ProDOL or LynG probes, were established through the Phoenix ampho retroviral transduction system utilising pBABE-ProDOL and pBABE-LynAnchor-eGFP (LynG) plasmids. These plasmids were introduced into the Phoenix-Ampho virus packaging cell line via calcium phosphate precipitation for 6 h, resulting in the formation of replication-deficient viral particles for subsequent transduction. Harvested virus was collected 24 h post-transfection after passing through 0.22 µm syringe filters.

The transduction of Huh-7.5^38^, H838 (NCI-H838, ATCC), or HeLa (ATCC) cells was achieved through spin transduction at 340× g for 3 hours. The selection process of transduced cells with puromycin was initiated 24 hours after transduction. For single-molecule imaging, Huh-7.5 and HeLa ProDOL and LynG cell lines were sorted by FACS to yield cell lines with suitable expression levels.

Stable transgenic Jurkat CD4 T cell lines, expressing ProDOL or LynG, were established using a second generation lentiviral system. HEK293T cells were transfected with lentiviral vectors VSV-G and PAX2 in combination with a pWPI vector (pWPI-ProDOL or pWPI-LynAnchor-eGFP (LynG) plasmid) at a 1:2:3 ratio using polyethyleneimine (1 µg/µL). 48 h post transfection, supernatant was collected and filtered using 0.48 µm filters.

3×10^6^ Jurkat CD4 T cells were resuspended in 1 ml viral supernatant and centrifuged at 1,000× g for 90 min. 5 ml media was added 16 h post transduction. For single-molecule imaging, Jurkat CD4 T cells were sorted through FACS to yield cell lines with suitable expression levels.

### FACS

Fluorescence activated cell sorting was performed on an Aria III cell sorter using FACSDiva version 8.0.1 (BD Biosciences). The gating of live cells was carried out based on the front and side scatter signal. Further sorting was conducted based on the fluorescence signal generated from 488 or 633 nm excitation for eGFP- and 1 nM Halo-SiR-labelled samples respectively. The sorted cells were gathered into tubes filled with pre-warmed complete growth medium supplemented with 100 U/mL penicillin and 100 µg/mL streptomycin. Expression of full-length ProDOL and LynG probes by the established cell-lines was subsequently confirmed by Western blot.

### Western blot

To examine the Nef-eGFP and eGFP expression cells were lysed in RIPA buffer while mixing on a rotator for 30 min before sonication on ice. The samples were then centrifugated at 20,000 g for 10 min and the supernatant was collected. Sample buffer (64 g/l SDS, 40 mM Tris pH=7.4, 8% glycerol, 12.3 g/l Dithiothreitol (DTT),, 0.16 g/l bromophenol blue, 20% 2-mercaptoethanol) was added and the mixture was heated for 2 min at 95 °C. Samples were run on a 10% sodium dodecyl sulphate polyacrylamide gel (SDS-PAGE) before blotting on a polyvinylidene fluoride (PVDF) membrane in Laemmli buffer for 1 h at 260 mA. Membranes were blocked in 5% BSA in tris-buffered saline 0.2% Tween-20 (TBST) before incubating overnight at 4 °C with primary anti-GFP, anti-His_6_ or anti-actin in 1% BSA in TBST. After washing with TBST the membranes were incubated for 1h with secondary antibodies conjugated with horseradish peroxidase at room temperature. Membranes were washed with TBST and developed in a 1:1 mixture of 0.1 M Tris at pH =8.5, 1.1 mg/l luminol, 0.185 mg/l p-coumaric acid, 1% DMSO, 0.018% H_2_O_2_.

### Cell Seeding

Lab-Tek 8-well chamber slides were cleaned by incubating twice with 0.1 M hydrofluoric acid for 30-60 s followed by washing twice with water. Thereafter Lab-Teks were incubated with PBS for at least 5 min. For adherent cell lines, chambers were used without further modification at a concentration of 1×10^4^cells/cm^2^.

For T cell experiments, 200 µl of 0.01% poly-L-lysine (Sigma-Aldrich) solution was added and incubated for 10 min before removing solution and letting it air-dry. Subsequently, the chambers were incubated for 3 h at 37 °C with 100 µl of α-CD3-antibody solution (10 µg/ml in PBS). Chambers were then washed 3 times with PBS before seeding of cells.

For ProDOL measurements, 2×10^4^ cells in 100 µL PBS were added to each chamber post labelling and incubated for 10 min before fixation with freshly prepared 4% paraformaldehyde (PFA). For time resolved activation studies 2×10^6^ cells in 100 µL PBS were added to each chamber and after 3 seconds fully aspirated. After full aspiration, 100 µL PBS was gently added and incubated for 1.5 min, 5 min, or 10 min before fixation with 4% PFA.

### Transfection

Transient transfection of adherent cells was carried out using FuGENE HD. 0.2 µg DNA, and 0.6 µL FuGENE HD are added to 10 µL OptiMEM. Mixture is incubated for 15 min at room temperature before adding to the cells in Lab-Tek 8-well chamber slide. 24 h post transfection, cells were used for further analysis.

Jurkat CD4 T cells were transiently transfected with eGFP or Nef-eGFP expression plasmids by electroporation. 5×10^6^ cells were collected by centrifugation, washed with 10 ml of serum-free media, resuspended in 500 µl serum-free media and transferred into a 0.4 cm electroporation vial. 20 µg of respective plasmid DNA was added, and the electroporation was performed at 250 V and 950 µF for 21 ms using the Gene Pulser XCell. Afterwards, the cells were transferred to a fresh 6-well chamber with 5 ml complete growth medium. 24 h post transfection, cells were used for further analysis.

### Labelling protocols

#### Pre- and post-fixation labelling of adherent cells

Cells expressing ProDOL probe or LynG were stained with 0.1–1,000 nM of Halo-dye or SNAP-dye conjugate for 0.25 -16 h in live or fixed cells. Cells were then washed four times with pre-warmed DMEM at 37 °C and 5% CO_2_ for 15 min, 15 min, 45 min and 5 min. Cells were fixed with freshly prepared and pre-warmed electron-microscopy-grade 3.7% PFA (EMS) in PBS for 40 min. All cells were imaged in serum-free media.

#### U2OS nuclear pore complex labelling

For cross validation analysis U2OS cells were fixed with freshly prepared 2.4% PFA for 30 seconds, before permeabilization with 0.4% (v/v) Triton X-100 in PBS, followed by fixation again in 2.4% PFA. Samples were quenched with 100 mM NH4Cl in PBS and washed twice for 5 min before a 30 min incubation in Image-iT Signal Enhancer. Samples were stained with 1 µM SNAP-AF647 diluted in 1 µM DTT, 0.5% BSA in PBS for 2 hours. The samples were washed 3 times in PBS for 5 min before imaging.

#### Jurkat CD4 T cell HaloTag staining

For live cell staining Jurkat CD4 T cells were collected by centrifugation for 3 minutes at 200× g after which the media was replaced with full growth RPMI-1640 media containing 0.1 nM to 100 nM Halo-SiR for 15 min to 16 hours at 37 °C (Fig. 3b), 10 nM for variable durations (Fig. 4a), or 1 nM for 1 h (Fig. 4c). The cells are then washed by spinning the sample down and replacing the staining solution with fresh media after 15 min three times and then a final 40 min wash in RPMI. The cells are spun down one final time before being resuspended in PBS.

#### Phospho-SLP-76

Antibody staining of Jurkat CD4 T cells was conducted on cells already fixed on the coated coverslips. The sample is permeabilised with 0.1% Triton X-100 (Sigma-Aldrich) with 1 mM Na3VO4 in PBS for 5 min before being washed three times with 1 mM Na3VO4 in PBS. The sample was blocked with PBS containing 5% FBS and 1 mM Na3VO4 for 30 min before incubating overnight with 2 μg/ml anti-pY145 SLP-76. Finally, the sample was washed three times with PBS before post fixation with 2% PFA in PBS for 15 min.

### Microscopy setup

#### Counting by Photon Statistics (CoPS)

CoPS measurements were performed on two custom-built confocal microscopes.

Microscope I (data acquired: Fig. 4 and Supplementary Fig. 5) (Axiovert 100, Zeiss) was equipped for sample-scanning confocal optical microscopy. The microscope setup included a XY & Z piezostage (Physik Instrumente) for nanometre resolution positioning and an AlphaPlan-Fluar 100×/1.45 oil immersion objective (Zeiss). The microscope included a <90 ps-pulsed laser diode emitting at 640 nm (LDH P-C-640B, PicoQuant, 20MHz repetition rate) and four single-photon sensitive avalanche photodiodes (APDs) (Perkin-Elmer). The excitation lasers were coupled into a single-mode polarisation maintaining fiber (Schäfter Kirchhof). A dichroic mirror (z532/640, CHROMA) was employed to separate the paths of the emission and excitation beams. The emitted signal was filtered using a quadband notch filter with additional spatial filtering using a 100 µm pinhole placed in the focal plane between two achromatic doublet lenses. All emission was split into four equal intensity paths using 50:50 beamsplitters and focused on the four APDs. 685/70 nm bandpass filters were place in front of each APD. Signals detected by the APDs were processed using a multichannel time-correlated single photon counting system (HydraHarp400, PicoQuant). The microscope was operated using SymPhoTime 64. The exposure settings, and the illumination intensities were tuned for each sample.

Microscope II (data acquired: Fig. 2 and Supplementary Fig. 6) (Eclipse Ti; Nikon) was equipped for laser-scanning confocal optical microscopy (Flimbee, PicoQuant). The microscope setup included a motorised XY-stage (Marzhäuser), a PFS2 autofocus system and an Apo TIRF 100× 1.49 NA oil immersion objective (Nikon). The microscope included a <90 ps-pulsed laser diode emitting at 640 nm (LDH P-C-640B, PicoQuant, 20MHz repetition rate) and four APDs (Perkin-Elmer). The excitation lasers were coupled into a single-mode polarisation maintaining fibre (Schäfter Kirchhof). A dichroic mirror (z532/640, CHROMA) was employed to separate the paths of the emission and excitation beams. The emitted signal was filtered using a quadband notch filter with additional spatial filtering using a 100 µm pinhole placed in the focal plane between two achromatic doublet lenses. All emission was split into four equal intensity paths using 50:50 beamsplitters and focused on the four APDs. 685/70 nm bandpass filters were place in front of each APD. Signals detected by the APDs were processed using a multichannel time-correlated single photon counting system (HydraHarp400, PicoQuant). The microscope was operated using SymPhoTime 64. The exposure settings, and the illumination intensities were each tuned for each individual sample.

#### Total internal reflection fluorescence (TIRF) microscope

ProDOL and quickPBSA measurements were carried out on a custom-built widefield microscope (Eclipse Ti; Nikon) equipped with both epifluorescence and total internal reflection fluorescence (TIRF) illumination. The microscope setup included a motorised XY-stage (Marzhäuser), a PFS2 autofocus system and an Apo TIRF 100× 1.49 NA oil immersion objective (both Nikon). Images were captured using an iXon Ultra 897 back-illuminated emCCD camera (Andor). The microscope utilised a fibre-coupled multi-laser engine (MLE; TOPTICA Photonics), equipped with 405 488, 561, and 640 nm laser lines for illumination. A quadband dichroic mirror was employed to separate the paths of the emission and excitation beams. The emitted signal was filtered using a quadband notch filter with additional bandpass filters (525/50, 605/70, and 690/70 nm), which were installed in a motorised filter wheel (Thorlabs) between body and an OptoSplit II (CAIRN). The microscope was operated using µManager 1.4^39^. The exposure times, the electron-multiplying gain, and the illumination intensities were each adjusted for each sample.

#### dSTORM super-resolution microscope

ELE measurements were carried out on a custom-built widefield microscope (RAMM, ASI) equipped with both epifluorescence and TIRF illumination. The microscope setup included a motorised XY-stage (ASI), a CRISP autofocus (ASI) and an Apo TIRF 100× 1.49 NA oil immersion objective (Nikon). Images were captured using a Prime95B sCMOS camera (Photometrics) with 130×130 µm field of view at 111.5 nm/px. The microscope utilised four laser lines for illumination: 405 nm (iBEAM-smart-405, Toptica); 488 nm (Cyan Laser-40, Spectra Physics); 561 nm (gem 561, Laser Quantum); 647 nm (2RU-VFL-P-2000-647, MPBC); controlled by two AOTFnC-Vis-TN (AA Opto-electronic) and passed through a beam shaper (piShaper, AdlOptica) for homogeneous illumination. A quadband dichroic mirror was employed to separate the paths of the emission and excitation beams. The emitted signal was filtered using a quadband notch filter with additional bandpass filters (450/40, 525/50, 593/46, and 731/137 nm), which were installed in a motorised filter slider (Thorlabs) between microscope body and camera. The microscope was operated using µManager 2.0^39^ extended with custom microcontroller boards (Arduino). Exposure times, gain, and illumination intensities were adjusted for each sample.

### Image acquisition

#### CoPS

Overview images were generated for each cell by confocal scanning. Protein clusters were localised using a custom-written analysis routine to detect local intensity maxima in acquired images. Spatially isolated maxima with a minimum distance of 500 nm to the next maxima position were selected for CoPS data acquisition. 1×10^7^ laser cycles (0.5 s) of photon coincidence events were recorded per cluster at 10 µW excitation power (measured before objective).

#### quickPBSA

Photobleaching step analysis was performed in ROXS PCD buffer^1^. Traces for U2OS Nup107-SNAP cells were acquired at an irradiance of 1.2 kW/cm^2^ at 640 nm. 3,000 frames with an exposure time of 200 ms per frame were recorded for each acquisition.

#### ELE

Nuclear pore complexes in dSTORM were acquired at 35 ms exposure time, 20,000 frames, HiLo illumination, at ∼4 kW/cm^2^ irradiance at the sample plane. The 405 nm laser irradiance was gradually increased over time using a predetermined exponential function (max. 30 W/cm^2^). 1 mL of 35 mM MEA glucose oxidase-catalase buffer according to Jimenez *et al*. was used^40^ in a custom-made airtight sample holder.

#### ProDOL

ProDOL measurements were acquired at 50 ms exposure time, 10.6 mW (640 nm), 3.4 mW (561 nm), 6.9 mW (488 nm) laser power (measured before the objective) and a gain of 100. 10 to 20 frames were recorded and averaged per measurement. Images were acquired beginning with 640 nm excitation, followed by 561 nm excitation and finally 488 nm excitation. 2 min of photobleaching at 561 nm (17 mW) was carried out before acquisition of the 488 nm excitation channel to reduce the potential of Förster energy transfer (FRET) between eGFP and red-shifted fluorophores excited at 561 nm.

#### Antibody DOL calibration

Phospho-SLP-76 antibodies were labelled with SiR-NHS (Spirochrome) and purified using Zeba spin desalting columns (40K MWCO, Thermo Fisher Scientific). DOL analysis was performed according to Equation (1) and yielded an ensemble DOL of 1.27 at 65% protein yield after purification.

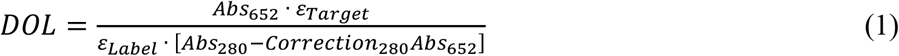

Using CoPS, the label number distribution per antibody could be generated (Supplementary Fig. 5a, b), yielding a pseudo-ensemble DOL of labelled antibodies of 1.38 with 80% of labelled antibodies carrying a single fluorophore, 9% carrying two and the remaining antibodies carrying >2 fluorophores. Using the measured distribution of fluorophores/antibody, the fraction of unlabelled antibodies could be determined to be 8%.

### Data analysis

#### ProDOL analysis

##### Segmentation

Binary masks to segment cells from background were generated from the reference channel as input with a custom-written imageJ script (processAverageIJwiththunderSTORM.ijm). In short, background was subtracted, high and low frequency domains were filtered from the image and a threshold was automatically generated. Finally, masks were exported as tiff files for subsequent analysis.

##### Emitter localisation

Single-molecule signals were localised with sub-pixel accuracy using ThunderSTORM called from a custom-written ImageJ script (processAverageIJwiththunderSTORM.ijm) with multi-emitter fitting enabled and a maximum number of 3 overlapping emitters. Localisations with sigma values outside a ±50% window around the modal PSF width were removed.

##### Registration

Registration of localisations in both target and reference channel were calculated in MATLAB (script: ProDOL_pipeline_thunderSTORM.m). One global transformation matrix for all images from the same condition was generated by registration of each localisation set. Localisation sets with <50 emitters were excluded. Additionally, all registrations with >3 pixels xy shift, >5° rotational shift and 5% scaling shift were excluded. All remaining transformation parameters were averaged and applied subsequently to all images in a given experiment.

##### Threshold determination

A distance cut-off T was calculated following Equation (2) in MATLAB (script: ProDOL_pipeline_thunderSTORM.m) for all image sets contained in an experiment. Colocalization distance cut-off *T* at which the fraction of specific colocalisation is maximised while the contribution of random colocalisation is kept at a minimum using Equation (2).

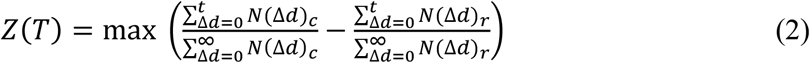

For each tolerance *t* a specific colocalisation score *Z* was determined at which the fraction of colocalising signals *N*_*c*_ is compared to a randomised colocalisation *N*_*r*_. Random colocalisation values are computed by rotating the tag target 90° relative to the reference. Δ*d* represents the distance of reference and target signal.

##### Density correction

Density correction was performed using Equation (3) (script: ProDOL_pipeline_thunderSTORM.m). Correction factors (CF_slope_ and CF_offset_) were determined from simulated data generated with testSTORM^41^. Simulated data was processed using the same approach as for experimental data described above and the recall as function of simulated emitter density was assessed to obtain parameters for CF_slope_ and CF_offset_ using linear regression (Supplementary Fig. 2).

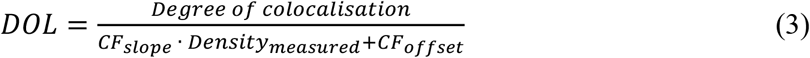

#### Classification

Manual classification of segmented data can be performed to remove image sets for which segmentation failed (script: imageSetInspector.m). Here, segmented mask and reference channel are displayed in random order to visually inspect segmentation results.

#### CoPS

A python library (pycops) was used to convert the raw.ptu data to .MBin files which contain multiple photon detections (mDEs) per laser pulse. The data from the first five to ten million laser pulses was then fitted to a mathematical model via a Levenberg-Marquardt algorithm. The data was collected and filtered for failed fits with a photon detection probability < 0.00075. For cellular DOL determination of Nup107-SNAP, the median number of detected emitters was divided by 32 (known stoichiometry).

#### quickPBSA

A custom Fiji script with the thunderSTORM plugin was used to determine the location of molecular clusters, after which the quickPBSA Python library was used to extract the intensity traces and correct for background intensity around each individual spot. The script also fitted the bleaching steps before summing up all the results. For cellular DOL determination of Nup107-SNAP, the median number of detected emitters was divided by 32 (known stoichiometry). Code availability: https://github.com/JohnDieSchere/quickpbsa.

#### ELE

All dSTORM data was analysed in SMAP^42^ using the fit_fastsimple workflow (dynamic factor: 1.3, with sCMOS correction and asymmetry enabled). Resulting localisations were analysed using the “SMAP_manual_NPC” manual related to Thevathasan *et al*. ^10^. Localisation were filtered (locprec: 0-12 nm, PSF: 110-185 nm, frame: 500-Inf; asymmetry: 0-0.2) and a NPC radius of 55 nm was assumed. NPC fitting was performed for all NPC with ≥3 labelled corners and a minimum localisation count of ≥2 emission events per corner.

#### Statistics

##### Curve fitting

All quantitative SLP-76 and phospho-SLP-76 distributions determined using CoPS (Fig. 4) were fitted using a log-normal distribution in GraphPad Prism v9.5.0 (730). SLP-76 does not form a cluster of consistent copy number and processes that involve a multiplicative product of independent random variables lead to a log-normal distribution (R^2^>0.85 for all fits).

Linear regression of detected fluorophores over degrees of labelling was calculated in GraphPad Prism using “simple linear regression” using mean, standard deviation (SD) and N for fluorophore numbers and mean, standard error (SE) for DOL measurements.

##### Statistical analysis

Comparison between DOL determination methods (Fig. 2) was performed using non-parametric ANOVA (Kruskal-Wallis) with outliers below 25% DOL removed from data analysis. In time series measurements (Fig. 4a), significance test of linear fits was determined using ANCOVA comparing the slope of linear fits to each other. phospho-SLP-76 histograms in the time series experiments were analysed using non-parametric ANOVA (Kruskal-Wallis). Levels of SLP-76 and phospho-SLP-76 in Nef experiments were compared to controls using non-parametric Kolmogorov-Smirnov. Significance testing of the phospho-SLP-76 to SLP-76-HaloTag ratio (Fig. 4) was determined using two-sample Z-test.

## Data availability

The data that support the findings of this study are available from the corresponding author upon reasonable request.

## Code availability

The ProDOL analysis package was written in the MATLAB and tested under version: 9.11.0 (R2021b). The ProDOL software is freely available under GNU General Public License v3.0 (https://github.com/hertenlab/ProDOL). Additional acquisition and analysis routines are available from the corresponding author upon reasonable request.

## Acknowledgements

We acknowledge funding from the Deutsche Forschungsgemeinschaft (DFG) through project PhotoQuant HE4559/6-1, by the Centre of Membrane Proteins and Receptors (COMPARE, Universities of Birmingham and Nottingham), and by the Academy of Medical Sciences (Grant APR2\1013). Oliver Fackler and Dirk-Peter Herten acknowledge shared funding by the Bundesministerium für Bildung und Forschung (BMBF/VDI, Switch-Click). Dirk-Peter Herten also acknowledges funding by the Bundesministerium für Bildung und Forschung (BMBF/VDI, LungSys). Some computations described in this paper were performed using the University of Birmingham’s BEAR Cloud service. We are grateful for the services provided by the FACS sorting facility at ZMBH (RI_00566) at Heidelberg University. We thank Ralf Bartenschlager for experimental support with the Huh-7.5 cell line. U2OS cells stably expressing Nup107-SNAP-tag were kindly provided by Jan Ellenberg (EMBL Heidelberg).

## Author contributions

J.E., S.A.T., and K.Y. performed all data analysis. J.E., S.A.T., K.Y., S.H., F.H., W.C., S.G., and Z.S. acquired the data. S.H. generated the ProDOL probe. S.H., F.H., K.Y. J.E., J.H. and S.A.T. generated the ProDOL analysis workflow. W.C., S.H., N.T., and F.S. generated stable cell lines. Conceptualization, supervision, resources, and project administration on T cell work was performed by O.T.F. and D-P.H.. O.T.F. and U.K. contributed general ideas and concepts. S.H. and D-P.H. conceived the method. J.E., S.A.T., K.Y., O.T.F., and D-P.H. wrote the manuscript with input from all authors.

