## Supplemental Information for "A General Method to Accurately Count Molecular Complexes and Determine the Degree of Labelling in Cells Using Protein Tags"

### Probe sequences

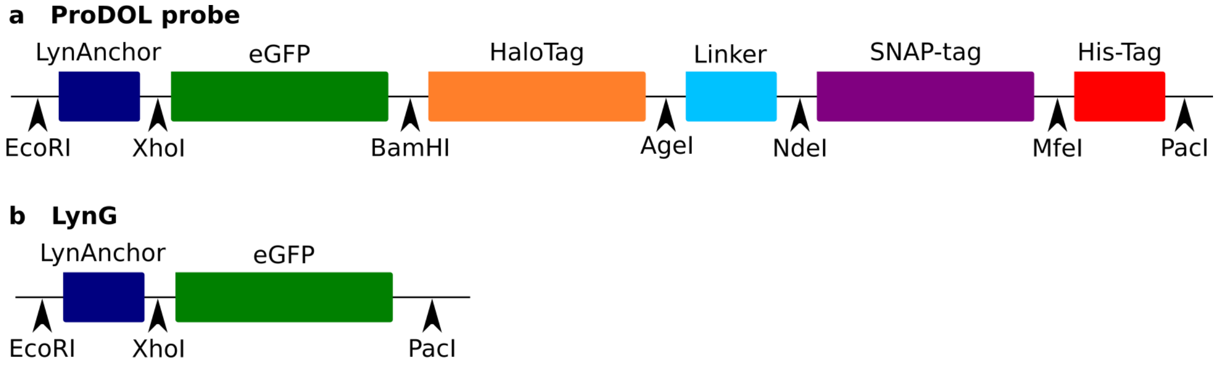


**Supplementary Fig. 1: ProDOL and LynG sequence features.** a, ProDOL consists of 6 domains (LynAnchor, eGFP, HaloTag, Linker, SNAP-tag, and His-Tag) separated by short (6 bp) unique restriction sites (EcoRI, XhoI, BamHI, AgeI, NdeI, MfeI, and PacI) allowing for easy modifiability of the construct. b, Truncated ProDOL probe named LynG (LynAnchor-eGFP) utilised as negative labelling control. For colour coded sequences see below.

Supplementary Information

**ProDOL**

ATGGGATGTATCAAGAGTAAGCGTAAGGATAATCTCAATGACGACGAGCTCGAGACCATGGTGAGCAAGGGCG
AGGAGCTGTTCACCGGGGTGGTGCCCATCCTGGTCGAGCTGGACGGCGACGTAAACGGCCACAAGTTCAGCGT
GTCCGGCGAGGGCGAGGGCGATGCCACCTACGGCAAGCTGACCCTGAAGTTCATCTGCACCACCGGCAAGCTG
CCCGTGCCCTGGCCCACCCTCGTGACCACCCTGACCTACGGCGTGCAGTGCTTCAGCCGCTACCCCGACCACATG
AAGCAGCACGACTTCTTCAAGTCCGCCATGCCCGAAGGCTACGTCCAGGAGCGCACCATCTTCTTCAAGGACGA
CGGCAACTACAAGACCCGCGCCGAGGTGAAGTTCGAGGGCGACACCCTGGTGAACCGCATCGAGCTGAAGGG
CATCGACTTCAAGGAGGACGGCAACATCCTGGGGCACAAGCTGGAGTACAACTACAACAGCCACAACGTCTATA
TCATGGCCGACAAGCAGAAGAACGGCATCAAGGTGAACTTCAAGATCCGCCACAACATCGAGGACGGCAGCGT
GCAGCTCGCCGACCACTACCAGCAGAACACCCCCATCGGCGACGGCCCCGTGCTGCTGCCCGACAACCACTACC
TGAGCACCCAGTCCAAACTGAGCAAAGACCCCAACGAGAAGCGCGATCACATGGTCCTGCTGGAGTTCGTGAC
CGCCGCCGGGATCACTCTCGGCATGGACGAGCTGTACAAGGGATCCGCAGAAATCGGTACTGGCTTTCCATTCG
ACCCCCATTATGTGGAAGTCCTGGGCGAGCGCATGCACTACGTCGATGTTGGTCCGCGCGATGGCACCCCTGTG
CTGTTCCTGCACGGTAACCCGACCTCCTCCTACGTGTGGCGCAACATCATCCCGCATGTTGCACCGACCCATCGCT
GCATTGCTCCAGACCTGATCGGTATGGGCAAATCCGACAAACCAGACCTGGGTTATTTCTTCGACGACCACGTCC
GCTTCATGGATGCCTTCATCGAAGCCCTGGGTCTGGAAGAGGTCGTCCTGGTCATTCACGACTGGGGCTCCGCT
CTGGGTTTCCACTGGGCCAAGCGCAATCCAGAGCGCGTCAAAGGTATTGCATTTATGGAGTTCATCCGCCCTATC
CCGACCTGGGACGAATGGCCAGAATTTGCCCGCGAGACCTTCCAGGCCTTCCGCACCACCGACGTCGGCCGCA
AGCTGATCATCGATCAGAACGTTTTTATCGAGGGTACGCTGCCGATGGGTGTCGTCCGCCCGCTGACTGAAGTC
GAGATGGACCATTACCGCGAGCCGTTCCTGAATCCTGTTGACCGCGAGCCACTGTGGCGCTTCCCAAACGAGCT
GCCAATCGCCGGTGAGCCAGCGAACATCGTCGCGCTGGTCGAAGAATACATGGACTGGCTGCACCAGTCCCCTG
TCCCGAAGCTGCTGTTCTGGGGCACCCCAGGCGTTCTGATCCCACCGGCCGAAGCCGCTCGCCTGGCCAAAAG
CCTGCCTAACTGCAAGGCTGTGGACATCGGCCCGGGTCTGAATCTGCTGCAAGAAGACAACCCGGACCTGATCG
GCAGCGAGATCGCGCGCTGGCTGTCGACGCTGGAGATTTCCGGCACCGGTTTGGCGGAGGCGGCGGCGAAG
GAGGCGGCGGCGAAGGAGGCGGCGGCGAAGGAGGCGGCGGCGAAGGCGGCGGCGCATATGGACAAAGACT
GCGAAATGAAGCGCACCACCCTGGATAGCCCTCTGGGCAAGCTGGAACTGTCTGGGTGCGAACAGGGCCTGCA
CCGTATCATCTTCCTGGGCAAAGGAACATCTGCCGCCGACGCCGTGGAAGTGCCTGCCCCAGCCGCCGTGCTGG
GCGGACCAGAGCCACTGATGCAGGCCACCGCCTGGCTCAACGCCTACTTTCACCAGCCTGAGGCCATCGAGGA
GTTCCCTGTGCCAGCCCTGCACCACCCAGTGTTCCAGCAGGAGAGCTTTACCCGCCAGGTGCTGTGGAAACTGC
TGAAAGTGGTGAAGTTCGGAGAGGTCATCAGCTACAGCCACCTGGCCGCCCTGGCCGGCAATCCCGCCGCCAC
CGCCGCCGTGAAAACCGCCCTGAGCGGAAATCCCGTGCCCATTCTGATCCCCTGCCACCGGGTGGTGCAGGGC
GACCTGGACGTGGGGGGCTACGAGGGCGGGCTCGCCGTGAAAGAGTGGCTGCTGGCCCACGAGGGCCACAG
ACTGGGCAAGCCTGGGCTG GGTCAATTGCACCATCACCATCACCAC

**LynG**

ATGGGATGTATCAAGAGTAAGCGTAAGGATAATCTCAATGACGACGAGCTCGAGACCATGGTGAGCAAGGGCG
AGGAGCTGTTCACCGGGGTGGTGCCCATCCTGGTCGAGCTGGACGGCGACGTAAACGGCCACAAGTTCAGCGT
GTCCGGCGAGGGCGAGGGCGATGCCACCTACGGCAAGCTGACCCTGAAGTTCATCTGCACCACCGGCAAGCTG
CCCGTGCCCTGGCCCACCCTCGTGACCACCCTGACCTACGGCGTGCAGTGCTTCAGCCGCTACCCCGACCACATG
AAGCAGCACGACTTCTTCAAGTCCGCCATGCCCGAAGGCTACGTCCAGGAGCGCACCATCTTCTTCAAGGACGA
CGGCAACTACAAGACCCGCGCCGAGGTGAAGTTCGAGGGCGACACCCTGGTGAACCGCATCGAGCTGAAGGG
CATCGACTTCAAGGAGGACGGCAACATCCTGGGGCACAAGCTGGAGTACAACTACAACAGCCACAACGTCTATA
TCATGGCCGACAAGCAGAAGAACGGCATCAAGGTGAACTTCAAGATCCGCCACAACATCGAGGACGGCAGCGT
GCAGCTCGCCGACCACTACCAGCAGAACACCCCCATCGGCGACGGCCCCGTGCTGCTGCCCGACAACCACTACC
TGAGCACCCAGTCCAAACTGAGCAAAGACCCCAACGAGAAGCGCGATCACATGGTCCTGCTGGAGTTCGTGAC
CGCCGCCGG GATCACTCTCGGCATGGACGAGCTGTACAAG

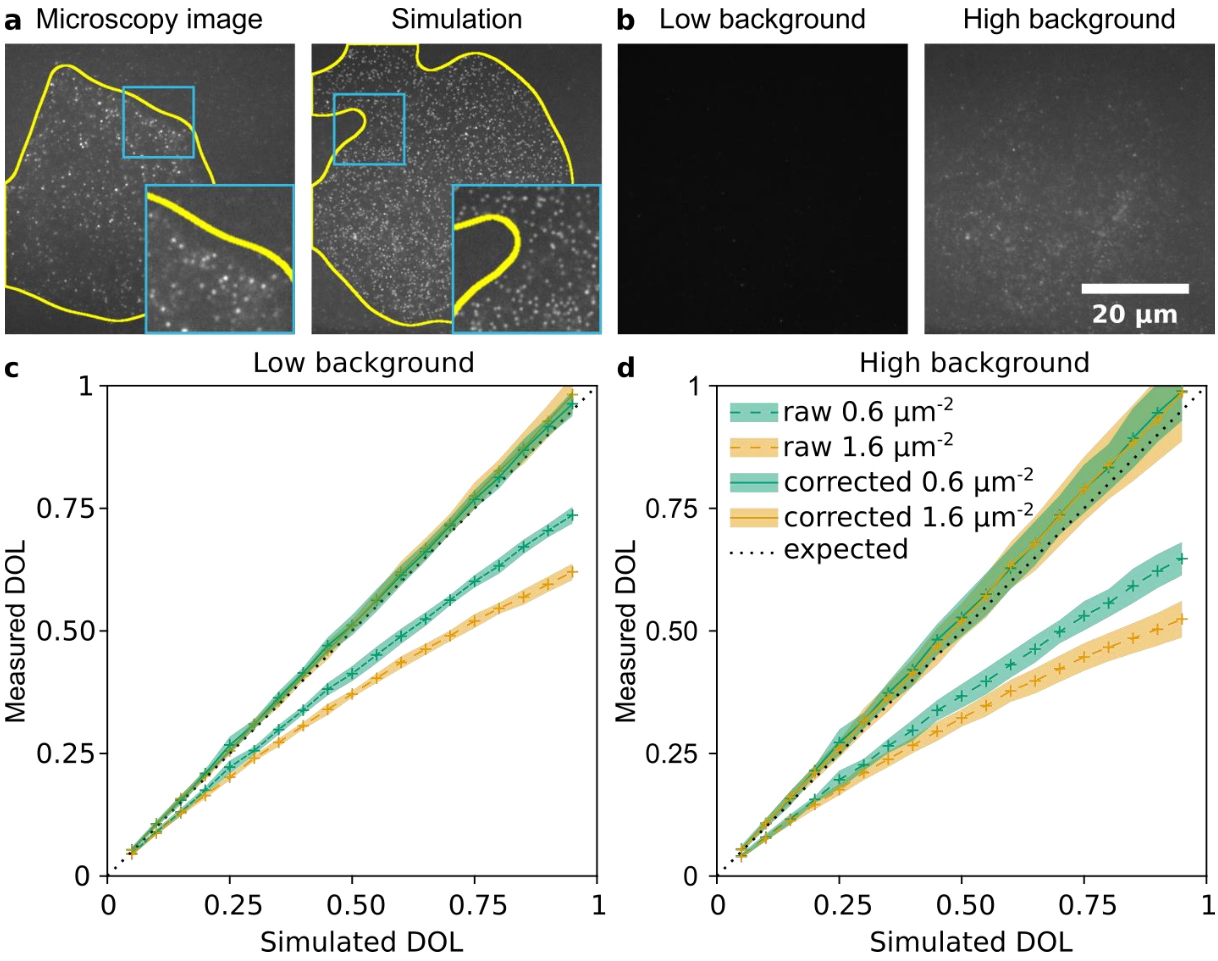


**Supplementary Fig. 2: ProDOL density correction.** a, Comparison of experimental (left) and simulated (right) microscopic data. Simulated images were generated by combining experimental data of cells expressing ProDOL without tag labelling and simulated emitters closely resembling experimental data at varying density and background. Yellow outline represents segmented cell. b, Example of high and low background from unlabelled H838 used in simulations. c,d, Simulated images were used to validate density corrections at reference density of 0.6 and 1.6 µm$^{-2}$ respectively (representative average emitter densities in cells). Median ± SD for raw data (dashed) and corrected (solid) DOL. n=20 per datapoint.

### Simulations for labelling efficiency measurements

Randomly distributed single emitter were simulated with testSTORM. A Gaussian PSF model with "Vesicles pattern" and acquisition parameters in line with experimental data was generated with drift disabled. “Alexa Fluor 647” dye model was adapted with the following parameters to resemble emitter properties matching experimental data: on time: 0.025, off time: 0.01, bleaching constant: 0.2, emitted photon/s: 350, number of labels per epitope: 1. Background was selected from a compilation of representative images of unlabelled cells imaged under identical acquisition settings as used for ProDOL. Low background images were acquired under TIRF illumination at 561 nm. High background images were acquired under TIRF at 640 nm excitation.

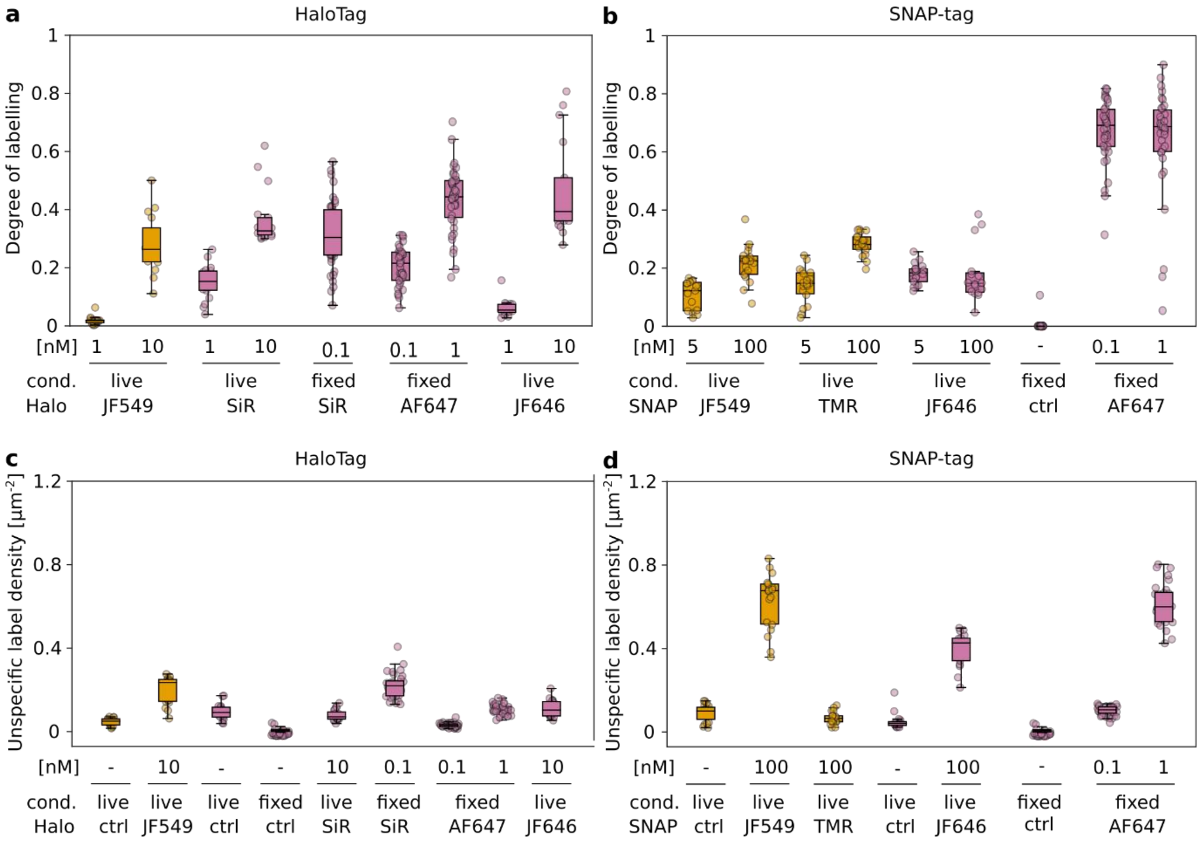


**Supplementary Fig. 3: Labelling efficiency and unspecific label density in Huh-7.5 cells.** Huh-7.5 cells expressing ProDOL for DOL determination (a,b) or LynG for unspecific label density determination (c,d). Cells were labelled for both HaloTag (a,c) and SNAP-tag (b,d) with varying dyes (live: 30 min, fixed: 120 min). Data represents cellular average, 15-42 cells per condition. Orange: 561 nm excitation, purple: 640 nm excitation.

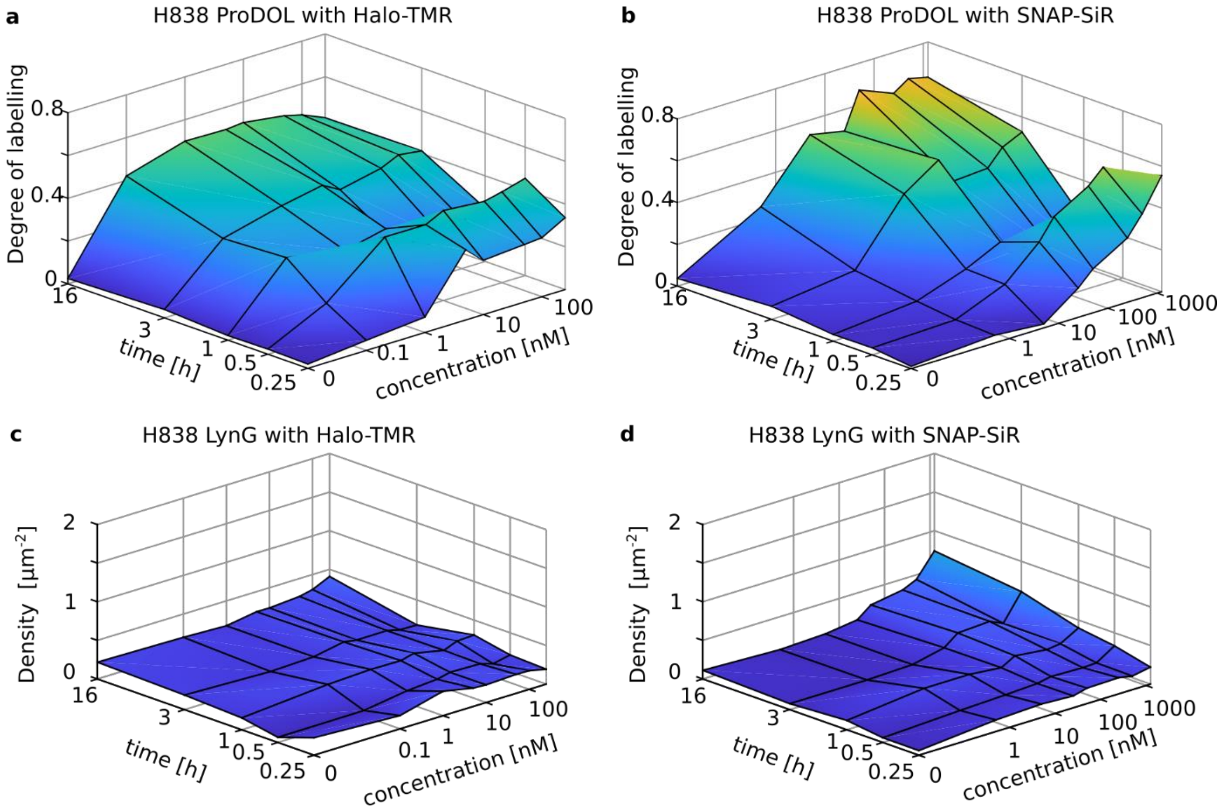


**e**

| | HaloTag with TMR | SNAP-tag with SiR |
|---|---|---|
| Cell type | $DOL_{max}$ [%] | $DOL_{max}$ [%] |
| H838 | 50 ± 12 | 67 ± 14 |
| HeLa | 42 ± 10 | 41 ± 5 |
| Huh7.5 | 42 ± 6 | 57 ± 14 |

**Supplementary Fig, 4: Determination of DOL as a parameter of incubation time and dye concentration. a**, H838 cells expressing ProDOL (a,b) and LynG (c,d) were stained with varying concentrations of Halo-TMR (a,c) and SNAP-SiR (b,d). a,b, Determined median DOL using ProDOL at varying ligand concentrations and incubation time. c,d, Unspecific labelling density measured at varying ligand concentrations and incubation time. e, Maximum DOL found for Halo-TMR and SNAP-SiR per cell line. Median from 4-15 cells per condition.

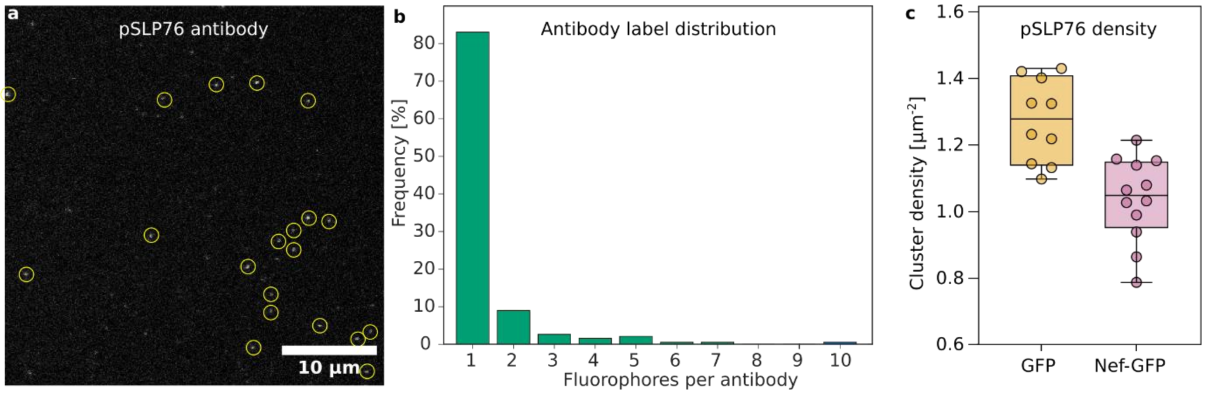


**Supplementary Fig. 5: Distribution of dyes per pSLP-76-antibody and pSLP-76 cluster density in cells.** a, representative microscopic image of immobilised pSLP-76 antibodies on a glass coverslip. b, Label distribution of pSLP-76 antibodies determined by CoPS. n=233. c, Distribution of cluster density of pSLP-76 in Jurkat cells expressing Nef-eGFP and negative control (eGFP).

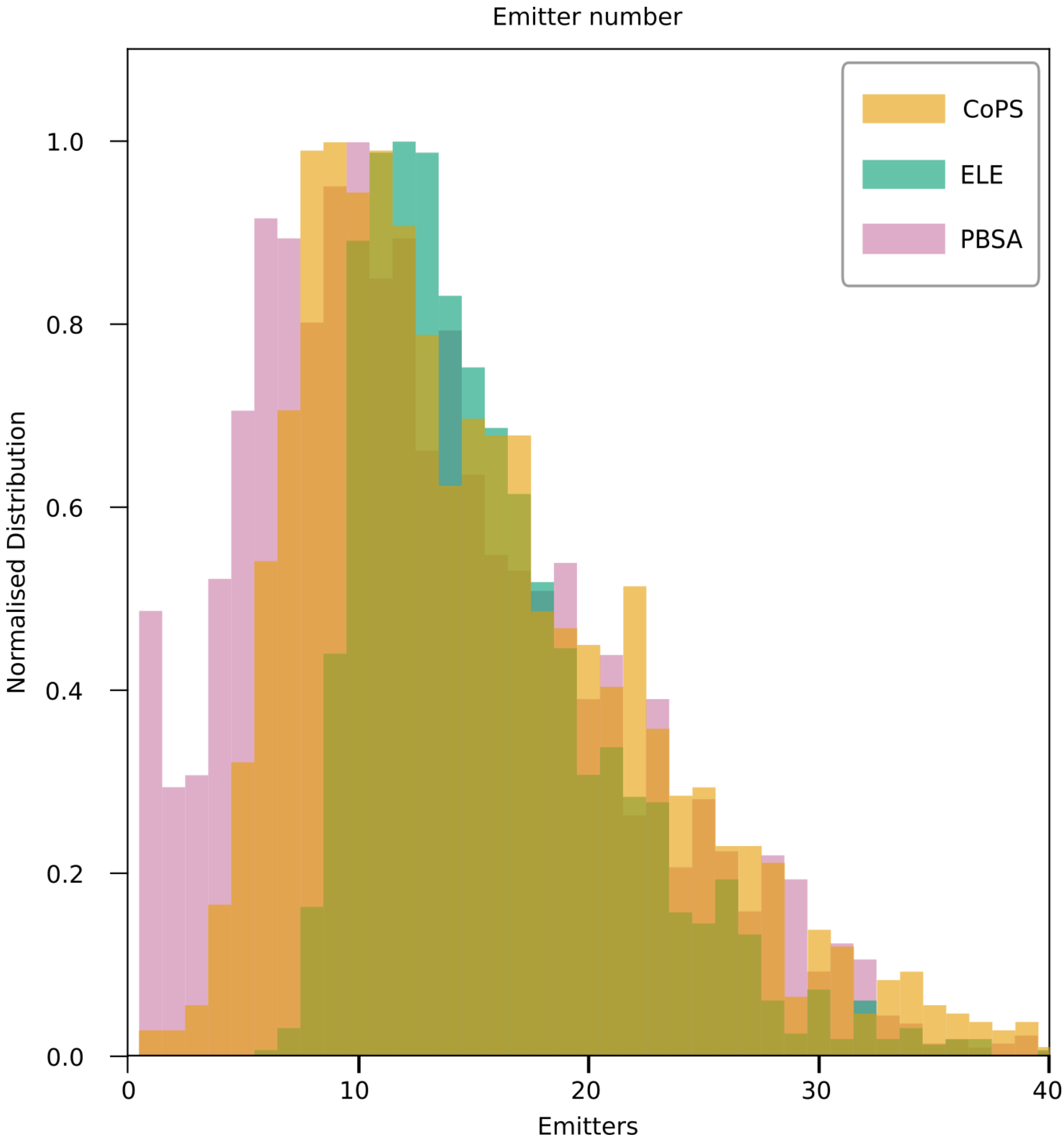


**Supplementary Fig. 6: Labelling distribution of Nup107-SNAP as found by CoPS, ELE and PBSA.** Data represents distribution of labels per individual nuclear pore complex (NPC) as determined by CoPS (orange), ELE (green), or PBSA (purple). ELE has a low detection rate at <8 emitters as a circular fit needs to be performed and therefore needs multiple corners labelled. ELE and PBSA both have detections of >32 labels, indicating counting of multiple NPC not detected in diffraction limited microscopy.
